# Experimental infection of elk (*Cervus canadensis*) and mule deer (*Odocoileus hemionus*) with SARS-CoV-2

**DOI:** 10.1101/2023.07.25.550568

**Authors:** Stephanie M. Porter, Airn E. Hartwig, Helle Bielefeldt-Ohmann, J. Jeffrey Root, Angela M. Bosco-Lauth

**Affiliations:** U.S. Department of Agriculture, Fort Collins, CO, USA; Colorado State University, Fort Collins, CO, USA; University of Queensland, St Lucia, Queensland, Australia

**Author notes:** **Address for Correspondence** Angela Bosco-Lauth, Department of Biomedical Sciences, Colorado State University, 1683 Campus Delivery, Fort Collins, CO 80523, USA,. These senior authors contributed equally to this article (J. Jeffrey Root, Angela M. Bosco-Lauth).

**Keywords:** cervid, *Cervus canadensis*, coronavirus, Delta variant, elk, mule deer, *Odocoileus hemionus*, SARS-CoV-2, wildlife

## Abstract

Elk (*Cervus canadensis*) and mule deer (*Odocoileus hemionus*) were experimentally evaluated for susceptibility to SARS-CoV-2. Elk did not shed infectious virus but produced low-level serological responses. Mule deer shed and transmitted virus in addition to mounting pronounced serological responses; they could therefore play a role in the epidemiology of SARS-CoV-2.

**Article Summary Line:** Experimental infection of elk (*Cervus canadensis*) and mule deer (*Odocoileus hemionus*) with SARS-CoV-2 revealed that while elk are minimally susceptible to infection, mule deer become infected, shed infectious virus, and can infect naïve contacts.

## Introduction

Many mammalian species from diverse taxonomies can be infected with SARS-CoV-2, as evidenced by both natural infections and experimental studies (1). Studies have identified the white-tailed deer ACE2 receptor as being closely homologous to that of humans. Among the cervids, sika deer (*Cervus nippon*), reindeer (*Rangifer tarandus*), and Père David’s deer (*Elaphurus davidianus*) are also predicted to be highly susceptible to SARS-CoV-2 based on ACE2 modeling (2-4). White-tailed deer are indeed susceptible to experimental infection with SARS-CoV-2, after which they shed infectious virus, and infected naïve conspecifics (5-7). Surveillance studies have demonstrated SARS-CoV-2 infection in free-ranging and captive white-tailed deer in the United States and Canada, as determined by the detection of viral RNA, antibodies to SARS-CoV-2, or virus isolation (8-11). Notably, the Alpha and Delta variants of concern have persisted in white-tailed deer populations following their displacement in humans (12, 13). As SARS-CoV-2 has likely repeatedly spilled over from humans into white-tailed deer and circulated within deer populations in North America, assessing the susceptibility of other cervid species is of great interest. Here, we assessed the susceptibility of elk (*Cervus canadensis*) and mule deer (*Odocoileus hemionus*) to the Delta variant of SARS-CoV-2.

## The Study

Six weanling elk (all female) and six yearling mule deer (five female, one male) were assessed for susceptibility to SARS-CoV-2. All animal work was approved by the Colorado State University Institutional Animal Care and Use Committee. Animals were procured from private vendors and group housed (3 individuals belonging to the same species per room) in an animal biosafety level-3 (ABSL-3) facility at Colorado State University. All animals were seronegative for SARS-CoV-2 prior to study commencement.

The Delta variant of SARS-CoV-2 (BEI Resources, NIAID, NIH: Isolate hCoV-19/USA/MD-HP05647/2021 (Lineage B.1.617.2)), was passaged a single time in Vero cells, then diluted in phosphate-buffered saline. Two animals per room were inoculated intranasally with 3.7-4.5 log_10_ plaque-forming units of virus as confirmed by back-titration of inoculum on Vero cells; the third animal in each room served as a direct contact.

Animals were assessed daily for attitude and signs of clinical disease, including lethargy, anorexia, nasal discharge, sneezing, coughing, and dyspnea. One mule deer (#3) was observed to be tachypneic and coughing with increased respiratory effort upon arrival, a clinical sign that continued throughout the weeklong acclimation period and until euthanasia at 3 dpi according to study schedule. As this respiratory pattern was present prior to the start of the study, we do not attribute it to SARS-CoV-2 infection. No other clinical signs were observed in any mule deer.

Animals were sedated with 50-100mg intramuscular xylazine for handling and sampling. Oral, nasal, and rectal swabs were collected on 0, 1, 2, 3, 5, 7, and 14 dpi. Plaque assays revealed that none of the elk shed infectious virus orally or nasally; elk rectal swabs were therefore not assessed. Reverse transcription PCR was performed on elk oral and nasal swabs collected through 7 dpi. SARS-CoV-2 RNA was recovered from either oral or nasal swabs collected between 1 and 5 dpi from three directly inoculated elk; all had cycle threshold values above 28 (Table 1). Plaque assay showed that three of the four directly inoculated mule deer shed infectious virus both orally and nasally. Both contact mule deer shed virus nasally, with one also shedding virus orally (Figures 1 and 2). Oral shedding of virus commenced on either 2 or 3 dpi and resolved by 7 dpi for directly inoculated mule deer, while the contact mule deer only had live virus detected from the oral swab collected on 7 dpi (Figure 1). Nasal shedding of virus was more staggered, with inoculated animals initially shedding on 1, 2, or 3 dpi, and continuing to have positive nasal swabs through 7 dpi (for the two direct inoculants remaining at that time point). Each of the contact mule deer only had a single positive nasal swab, collected on 3 or 7 dpi (Figure 2). Infectious virus was not recovered from any of the mule deer rectal swabs.

**Table 1.**
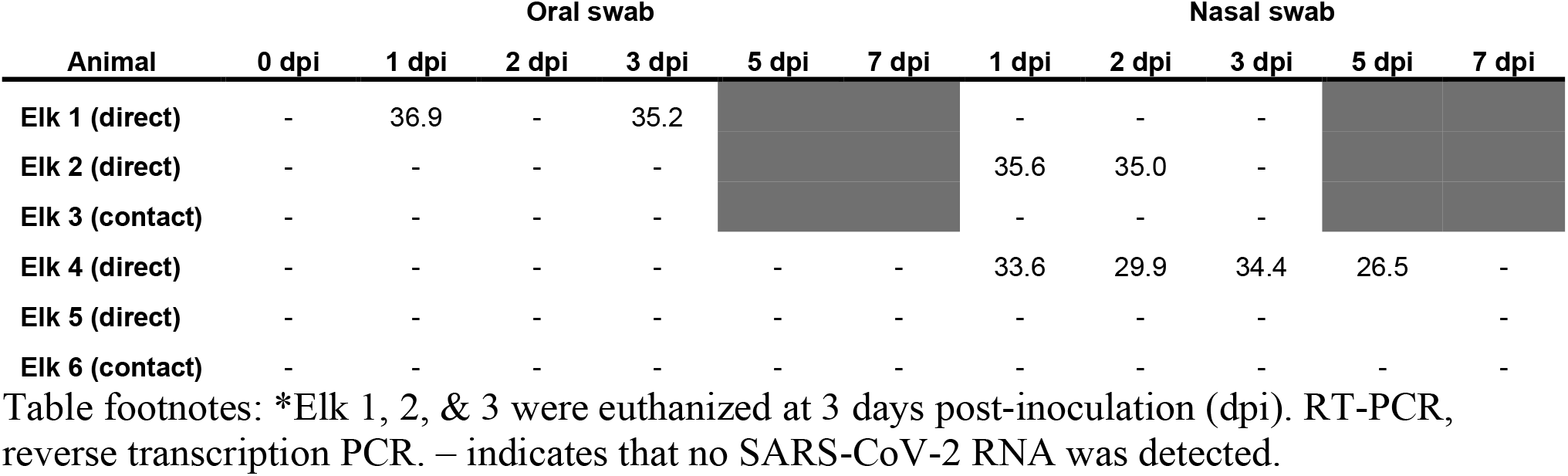
Cycle threshold values (RT-PCR) of oral and nasal swab samples from elk experimentally infected with SARS-CoV-2.

| Animal | Oral swab |  |  |  |  |  | Nasal swab |  |  |  |  |
| --- | --- | --- | --- | --- | --- | --- | --- | --- | --- | --- | --- |
|  | 0 dpi | 1 dpi | 2 dpi | 3 dpi | 5 dpi | 7 dpi | 1 dpi | 2 dpi | 3 dpi | 5 dpi | 7 dpi |
| Elk 1 (direct) | - | 36.9 | - | 35.2 |  |  | - | - | - |  |  |
| Elk 2 (direct) | - | - | - | - |  |  | 35.6 | 35.0 | - |  |  |
| Elk 3 (contact) | - | - | - | - |  |  | - | - | - |  |  |
| Elk 4 (direct) | - | - | - | - | - | - | 33.6 | 29.9 | 34.4 | 26.5 | - |
| Elk 5 (direct) | - | - | - | - | - | - | - | - | - |  | - |
| Elk 6 (contact) | - | - | - | - | - | - | - | - | - | - | - |

Table footnotes: \*Elk 1, 2, & 3 were euthanized at 3 days post-inoculation (dpi). RT-PCR, reverse transcription PCR. – indicates that no SARS-CoV-2 RNA was detected.

**Figure 1.**
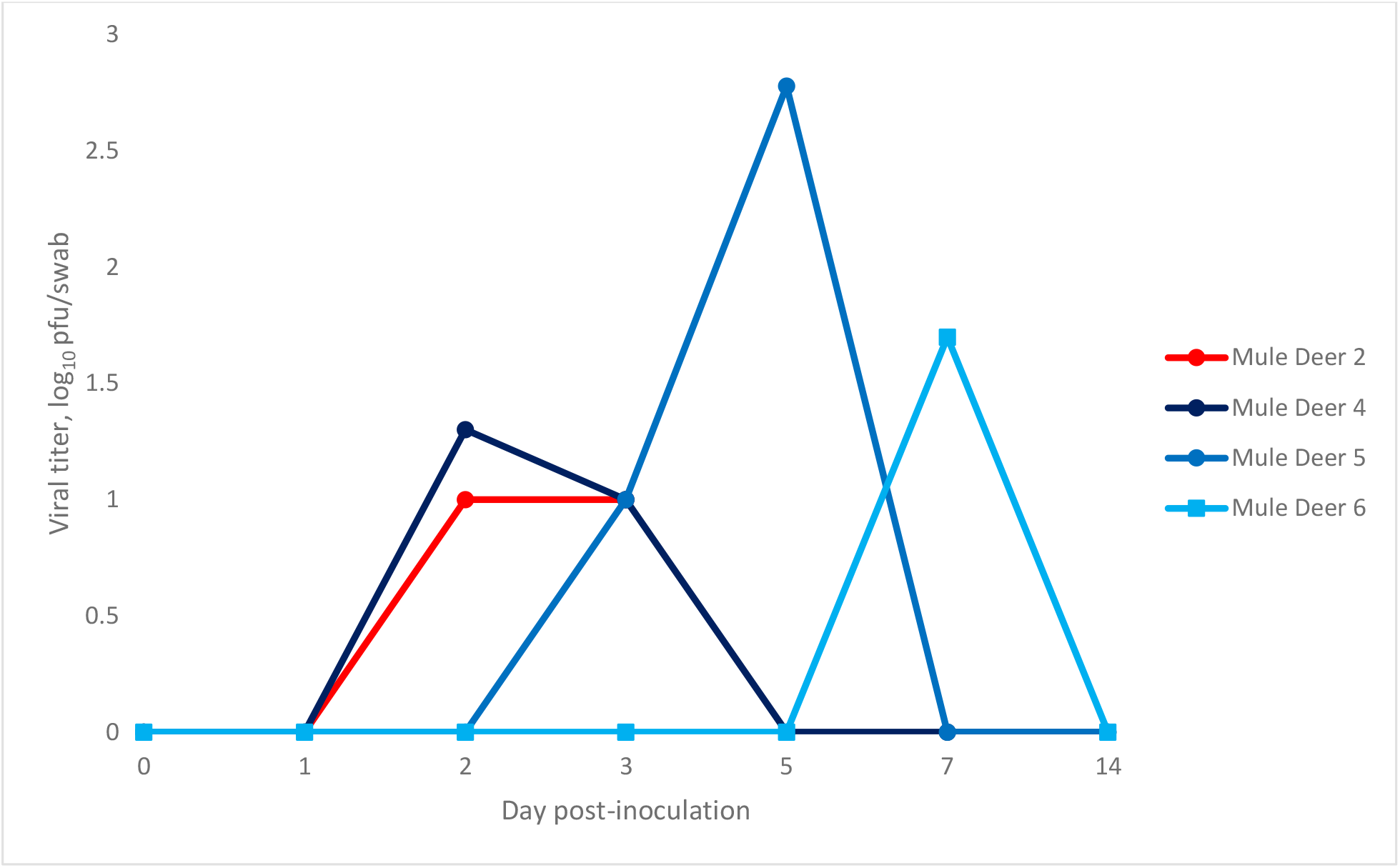
Oropharyngeal shedding of SARS-CoV-2 by mule deer as detected by plaque assay. Values are expressed as log_10_ plaque-forming units (pfu)/swab. Mule deer 2 was euthanized at 3 days post-infection (dpi). Mule deer 2, 4, and 5 were directly inoculated, while mule deer 6 was a contact animal.

**Figure 2.**
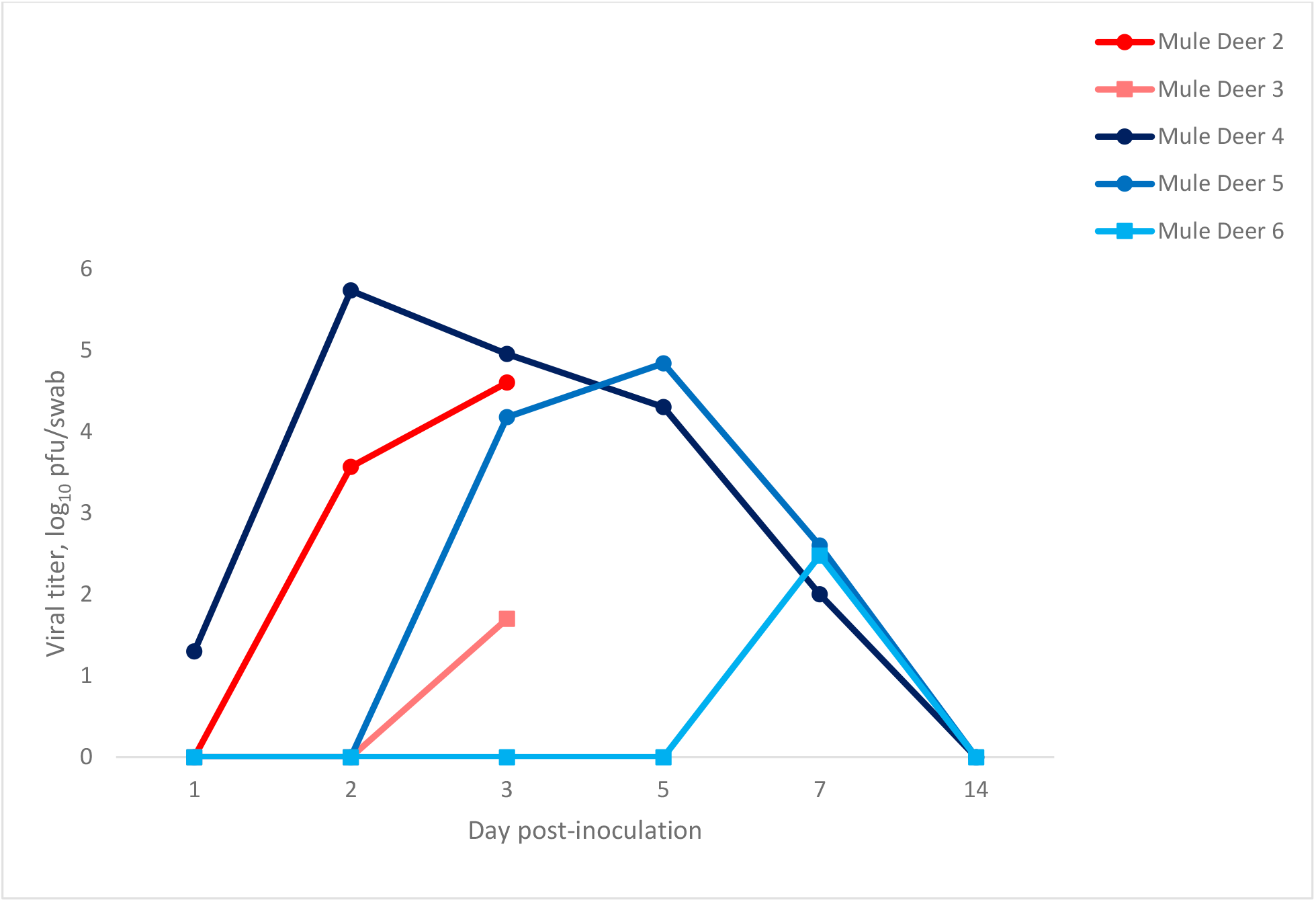
Nasal shedding of SARS-CoV-2 by mule deer as detected by plaque assay. Values are expressed as log_10_ plaque-forming units (pfu)/mL. Mule deer 2 and 3 were euthanized at 3 days post-infection (dpi). Mule deer 2, 4, and 5 were directly inoculated, while mule deer 3 and 6 were contact animals.

One room of each species (n=3, two inoculants and one contact) was humanely euthanized and necropsied on 3 dpi; tissues (nasal turbinates, trachea, heart, lung, liver, spleen, kidney, and small intestine) were collected for virus isolation and histopathology. While infectious virus was not detected in any of the elk tissues, infectious virus was recovered from the nasal turbinates and trachea of one directly inoculated mule deer.

The remaining animals were maintained until 21 dpi, at which point they were humanely euthanized, necropsied, and the aforementioned tissues collected into formalin. Blood from these animals was collected weekly and evaluated for a serological response to SARS-CoV-2 by plaque reduction neutralization test (14). Both directly inoculated elk developed a low-level antibody response, with peak neutralizing titers of 1:20 at 21 dpi. The contact elk did not seroconvert. All mule deer held until 21 dpi developed neutralizing antibody responses with peak titers reaching 1:1280 or higher (Table 2).

**Table 2.** Antibody titers (PRNT90) for elk and mule deer experimentally infected with SARS-CoV-2.

| Animal | Preinfection | 7 dpi | 14 dpi | 21 dpi |
| --- | --- | --- | --- | --- |
| Elk 4 | <10 | <10 | 10 | 20 |
| Elk 5 | <10 | <10 | 10 | 20 |
| Elk 6 | <10 | <10 | <10 | <10 |
| Mule deer 4 | <20 | 40 | 5120 | 1280 |
| Mule deer 6 | <20 | <20 | 1280 | 1280 |
| Mule deer 5 | <20 | <20 | 1280 | 640 |

Table footnotes: \*dpi = days post-inoculation

No gross lesions were appreciated in any animals at necropsy. Respiratory tissues from all mule deer were evaluated by a veterinary pathologist. The trachea of all mule deer were histologically unremarkable, while multifocal accumulations of mononuclear leukocytes in the absence of frank inflammation were noted in the nasal turbinates and/or lungs from five mule deer. Elk tissues were not evaluated histologically.

## Conclusions

Should wildlife populations serve as maintenance hosts for SARS-CoV-2, there could be substantial implications. For example, the persistence of viral variants that have been displaced in the human population, viral evolution, and spillback into a human, have all been suggested to have occurred in white-tailed deer populations (11, 12), although it is still unclear if the white-tailed deer will serve as maintenance hosts of the virus. Evaluating the susceptibility of other cervid species to SARS-CoV-2 will help direct surveillance efforts among free-ranging wildlife, which is key to both understanding the epidemiology of the virus and implementing control measures.

We challenged the animals in this experiment with the Delta variant of SARS-CoV-2 based on evidence that this variant of concern was prevalent in white-tailed deer populations (13). Our results indicate that while elk appear to be minimally susceptible to infection with the Delta variant of SARS-CoV-2, mule deer are highly susceptible and capable of transmitting the virus onwards. Infection in mule deer was subclinical, and while immune activation in the absence of frank inflammation was observed in the respiratory tissues from four of six animals, this finding may or may not be linked to SARS-CoV-2 infection. Notably, all mule deer used in this study were incidentally tested to assess their chronic wasting disease (CWD) status, and #1 and #3 were CWD positive. We do not believe that a concurrent infection with CWD greatly impacted their susceptibility to infection with SARS-CoV-2, as one of these animals became infected with SARS-CoV-2 and the other was the sole mule deer in this study which did not. Wild mule deer regularly become infected with CWD, therefore our study mimics the infection status in a natural setting (15).

While mule deer have a smaller distribution than white-tailed deer, they still represent an important population of cervids which is frequently in contact with humans and domestic animals. Therefore, susceptibility of mule deer provides yet another potential zoonotic source for SARS-CoV-2 spillover and/or spillback. At this time, there is no evidence that wildlife pose a significant source of SARS-CoV-2 exposure for humans, but the potential for this virus to become established in novel host species could lead to viral evolution in which novel variants may arise. Therefore, continued monitoring and surveillance of “at risk” species, such as white-tailed and mule deer, is necessary to detect any variants quickly and prevent onward transmission.

## Acknowledgements

This work was supported by internal funding from Colorado State University and the U.S. Department of Agriculture, Animal and Plant Health Inspection Service. This study was directly funded by the American Rescue Plan (or ARP) Act provision to conduct monitoring and surveillance of susceptible animals for SARS-CoV-2. The authors would like to thank and recognize the USDA APHIS Science Fellows Program for supporting the salary of SMP. We are very grateful to Tracy Nichols, Colorado and Kansas state veterinarians, and Colorado Parks and Wildlife for logistical support. Special thanks to Jeremy Ellis for technical assistance, Susan Shriner for consulting, McKinzee Barker, Elizabeth Lawrence, and Hannah Sueper for their help with animal handling, and to Richard Bowen for facilities support.

## Biographical sketch

Dr. Stephanie Porter is a Science Fellow with the National Wildlife Research Center at the United States Department of Agriculture. Her research interests include the pathogenesis and transmission of infectious pathogens.

